# Gene model for the ortholog of *Ilp5* in *Drosophila erecta*

**DOI:** 10.1101/2025.08.05.668546

**Authors:** Leon F. Laskowski, Bethany C. Lieser, Sarah L. Hewitt, Miryam Perez, Brittany E. Saciccioli, Jacqueline Wittke-Thompson, Scott Tanner, Rachel Sterne-Marr

## Abstract

Gene model for the ortholog of *Insulin-like peptide 5* (*Ilp5*) in the Dere_CAF1 Genome Assembly (GenBank Accession: GCA_000005135.1) of *Drosophila erecta*. This ortholog was characterized as part of a developing dataset to study the evolution of the insulin/insulin-like growth factor signaling pathway across the genus *Drosophila* using the Genomics Education Partnership gene annotation protocol for Course-based Undergraduate Research Experiences.

## Introduction

*This article reports a predicted gene model generated by undergraduate work using a structured gene model annotation protocol defined by the Genomics Education Partnership (GEP; thegep*.*org) for Course-based Undergraduate Research Experience (CURE)* (Rele *et al*., 2023). *The following information in quotes was mostly repeated from inaugural articles submitted by participants using the same GEP CURE protocol for annotating Drosophila species orthologs of Drosophila melanogaster genes in the insulin signaling pathway*.

“In this GEP CURE protocol students use web-based tools to manually annotate genes in non-model Drosophila species based on orthology to genes in the well-annotated model organism, the fruit fly *Drosophila melanogaster*. This allows undergraduates to participate in course-based research by generating manual annotations of genes in non-model species (Rele *et al*., 2023). Computational-based gene predictions in any organism are often improved by careful manual annotation and curation, allowing for more accurate analyses of gene and genome evolution (Mudge and Harrow, 2016; Tello-Ruiz *et al*., 2019). These models of orthologous genes across species, such as the one presented here, then provide a reliable basis for further evolutionary genomic analyses when made available to the scientific community.” (Myers *et al*., 2024).

We propose a gene model for the *D. erecta* (taxid 7220) ortholog of the *D. melanogaster* Insulin-like peptide 5 (*Ilp5*) gene. The genomic region of the ortholog corresponds to the uncharacterized protein LOC6546022 (RefSeq accession XP_001972234.2) in the Dere_CAF1 Genome Assembly of *D. erecta* (GenBank Accession: GCA_000005135.1; Drosophila 12 Genomes Consortium, 2007). This model is based on RNA-Seq data (Chirn *et al*., 2015; Ma *et al*., 2018) from *D. erecta* (PRJNA264407; PRJNA414017) and *Ilp5* in *D. melanogaster* using FlyBase release FB2022_04 (GCA_000001215.4; Larkin et al., 2021; Gramates *et al*., 2022; Jenkins *et al*., 2022).

“In animals and in flies, the insulin/insulin-like peptide (Ilp) signaling (IIS) pathway plays a central role in metabolism, development, and reproduction (Semaniuk *et al*., 2021). Signaling is initiated when insulin or insulin-like peptides, of which there are eight in *Drosophila melanogaster*, bind to their receptors. Ilp5 shows particular conservation from *Drosophila* to mammals, and was shown to lower blood glucose levels when administered to rats (Sajid *et al*., 2011). In *Drosophila melanogaster*, Ilp5 expression is highest in the insulin-producing cells in the adult brain, but also expressed in the gut, ovaries, and neuroendocrine cells of larvae (Semaniuk *et al*., 2021). Flies lacking Ilp5 function show defects in fertility (Grönke *et al*. 2010; Strilbystka *et al*., 2020), behavior (Semaniuk *et al*. 2018; Strilbystka *et al*., 2020), metabolism (Semaniuk *et al*. 2018; Strilbystka *et al*. 2020), and lifespan (Grönke *et al*., 2010).” (Lawson *et al*., 2024).

*“D. erecta* is part of the *melanogaster* species group within the subgenus *Sophophora* of the genus *Drosophila* (Sturtevant, 1939; Bock and Wheeler, 1972). It was first described by Tsacas and Lachaise (1974). *D. erecta* is found in west central Africa (https://www.taxodros.uzh.ch, accessed 1 June 2024; Markow and O’Grady, 2006) where it is found to breed primarily on the fruits of *Pandanus candelabrum*, a spiny evergreen shrub (Unwin, 1920; Lachaise and Tsacas, 1983).” Lieser et al., submitted.

## Results

### Synteny

The target gene, *Ilp5*, occurs on chromosome 3L in *D. melanogaster* and is nested within *CG43897*, flanked upstream by *Cyclin-dependent kinase 8* (*Cdk8*) and *Inhibitor-2* (*I-2*), and flanked downstream by *Insulin-like peptide 4* (*Ilp4*), *Insulin-like peptide 3* (*Ilp3*), *Insulin-like peptide 2* (*Ilp2*), which are all nested within *CG32052*, and *Insulin-like peptide 1* (*Ilp1*) (Figure 1A). The *tblastn* search of *D. melanogaster* Ilp5-PB (query) against the *D. erecta* (GenBank Accession: GCA_000005135.1) Genome Assembly (Drosophila 12 Genomes Consortium, 2007; Zimin *et al*., 2008) placed the putative ortholog of *Ilp5* within scaffold scaffold_4784 (CH954178.1) at locus LOC6546022 (XP_001972234.2) with an E-value of 9e-14 and a percent identity of 59.3 Furthermore, the putative ortholog is nested by LOC6546021 (XP_026833897.1) and flanked upstream by LOC6546008 (XP_001972237.1) and LOC6546023 (XP_015013220.1), which correspond to *CG43897, I-2*, and *Cdk8* in *D. melanogaster* (E-value: 0.0, 0.0, and 3e-158, respectively, and identity: 90.7%, 98.5%, and 89.1%, respectively, as determined by *blastp*; Figure 1, Altschul *et al*., 1990).

**Figure 1.**
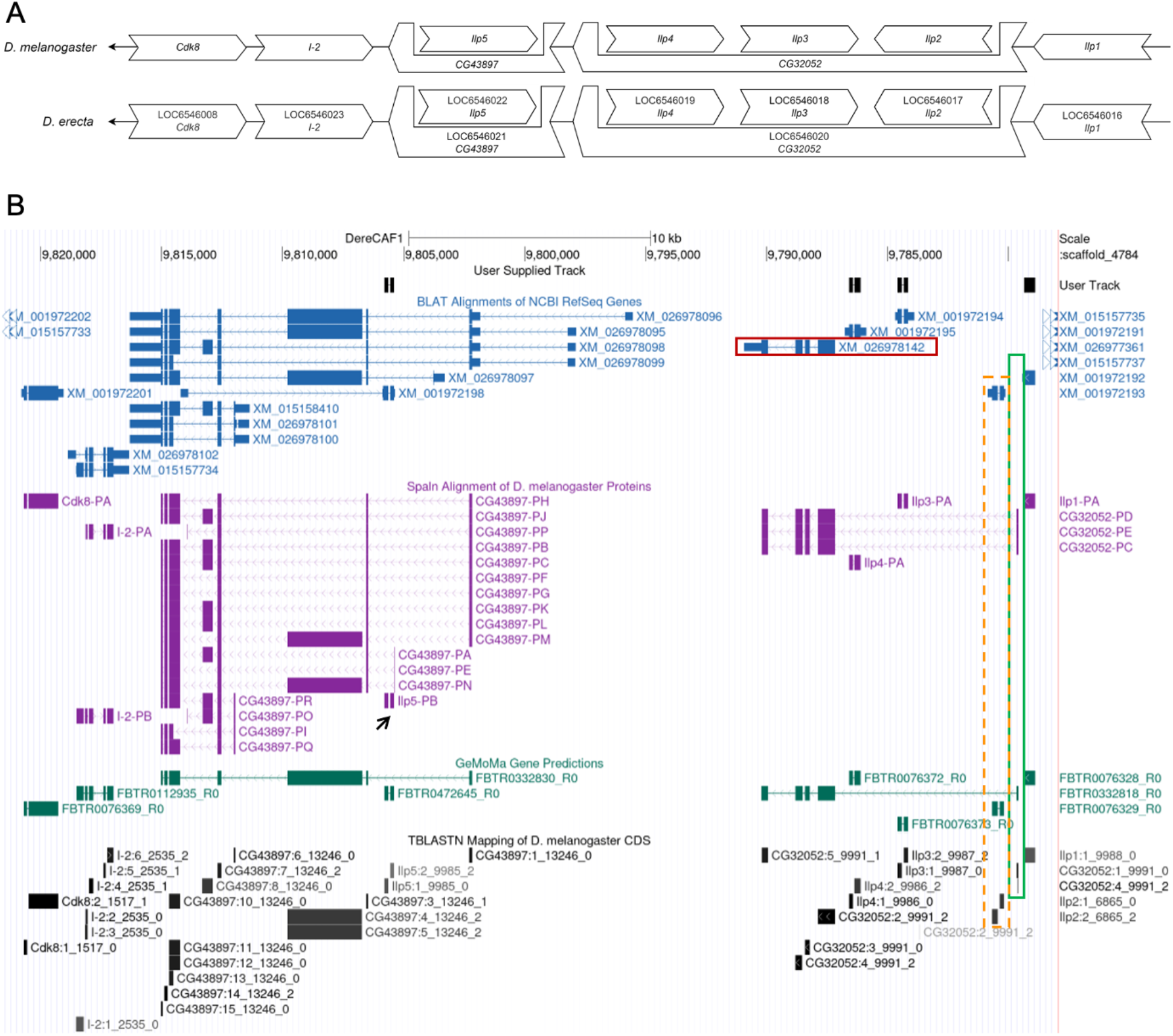
Synteny between *Drosophila erecta* and *Drosophila melanogaster* Genomic Neighborhoods. **A. Schematic Synteny Diagram**. Open/wide arrows indicate genes and loci (*D. erecta*). *Ilp5* genes in both species are located on the negative strand. The chromosome/scaffold direction has been reversed for both *D. melanogaster* and *D. erecta* in this diagram as indicated by the thin/closed arrows on the left pointing leftward. Gene symbols given in *D. erecta* indicate orthology to the *D. melanogaster genes*. Gene lengths and position are not drawn to scale. **B. *D. erecta* Genomic neighborhood of *Ilp5* shown in GEP UCSC Genome Browser Track Data Hub** (Raney *et al*., 2014). *Ilp5* (black arrow) is nested in *CG43897* downstream of *Cdk8* and *I-2. CG32052, Ilp4, Ilp3, Ilp2*, and *Ilp1* are downstream of *Ilp5*. The *CG32052* (LOC6546020) ortholog in *D. erecta* (red box) does not appear to be nesting *Ilp2, Ilp3*, and *Ilp4* as in *D. melanogaster* because the first CDS is not detected (green box) in the BLAT alignment (blue). TBLASTN mapping of *D. melanogaster* CDS track (black and gray) localizes exon 1_9991_0 to the same site in *D. erecta* as it does in *D. melanogaster*. The presence of the first CDS of *CG32052* in *D. erecta* is further supported by the GeMoMa Predictions (shown in dark green). *Ilp2* in *D. erecta* is not predicted by the Spaln alignment but is supported by TBLASTN mapping and GeMoMa Predictions (orange dashed box).

The putative ortholog of *Ilp5* is flanked downstream by LOC6546019 (XP_001972231.1), LOC6546018 (XP_001972230.1), LOC6546017 (XP_001972229.1), which are likely nested within LOC6546020 (XP_026833943.1), and LOC6546016 (XP_001972228.1), which correspond to *Ilp4, Ilp3, Ilp2, CG32052*, and *Ilp1* in *D. melanogaster* (E-value: 1e-62, 5e-65, 7e-90, 0.0, and 9e-79; identity: 84.1%, 92.6%, 91.2%, 98.3%, and 79.4%, respectively, as determined by *blastp*). *CG32052* in *D. melanogaster* nests *Ilp4, Ilp3*, and *Ilp2* within its first intron. However, in *D. erecta* the BLAT Alignment for the *CG32052* (LOC6546020) ortholog does not predict it to be a nesting gene. *CG32052* has five coding sequences (CDSs) in *D. melanogaster*, whereas *CG32052* is predicted to have only four CDSs in *D. erecta*, due to the absence of CDS1 (1_9991_0) (Figure 1B, red box). Although the BLAT alignment does not show *CG32052* nesting *Ilp4, Ilp3* and *Ilp2*, the CDS TBLASTN Mapping and GeMoMa Predictions support the presence of the CDS1 implying that those three genes in *D. erecta* are nested in *CG32052* (Figure 1B, Lieser *et al*., 2024). Therefore, we conclude that the structure of the *CG32052* gene in *D. erecta* is conserved relative to *D. melanogaster* and *Ilp4, Ilp3* and *Ilp2* are nested within *CG32052*.

The genes surrounding the *Ilp5* are orthologous to the genes at the *Ilp5* locus in *D. melanogaster* and synteny is likely completely conserved. Identifications of genes are supported by e-values and percent identities; thus, we conclude that LOC6546022 is the correct ortholog of *Ilp5* in *D. erecta* (Figure 1).

### Protein Model for *D. erecta* Ilp5

*Ilp5* in *D. erecta* has one protein-coding isoform (Ilp5-PB; Figure 2A) and the mRNA isoform contains two coding sequences (CDS). Both features are conserved with respect to the ortholog in *D. melanogaster*. The BLAT alignment and Geneid gene prediction suggest the use of nucleotides 9,805,847-9,805,845 as the ATG start codon (yellow box in Figure 2A). However, these predictions are not consistent with the RNA-Seq, the GeMoMa gene prediction or importantly the Spaln alignment of *D. melanogaster* Ilp5-PB. The GeMoMa prediction and Spaln alignment suggest that nucleotides 9,805,796-9,805,794 serve as the start codon (brown box in Figure 2A). The ortholog of Ilp5-PB is 108 amino acids in *D. melanogaster*, while the protein product in *D. erecta* is 110 amino acids due to a two amino acid insertion in the second CDS. The *blastp* search result shows 80% identity between *D. melanogaster* Ilp5-PB and the *D. erecta* gene model (E-value: 6e-54). Using word size of 3 to generate the dot plot results in several gaps along the diagonal (white boxes I, II, III, IV highlight largest; Figure 2B). However, the *blastp* search result indicates 89.1% similarity as also illustrated in the protein alignment (Figure 2C). The alignment further highlights that all but one amino acid in the first CDS are identical or similar between the two species.

**Figure 2.**
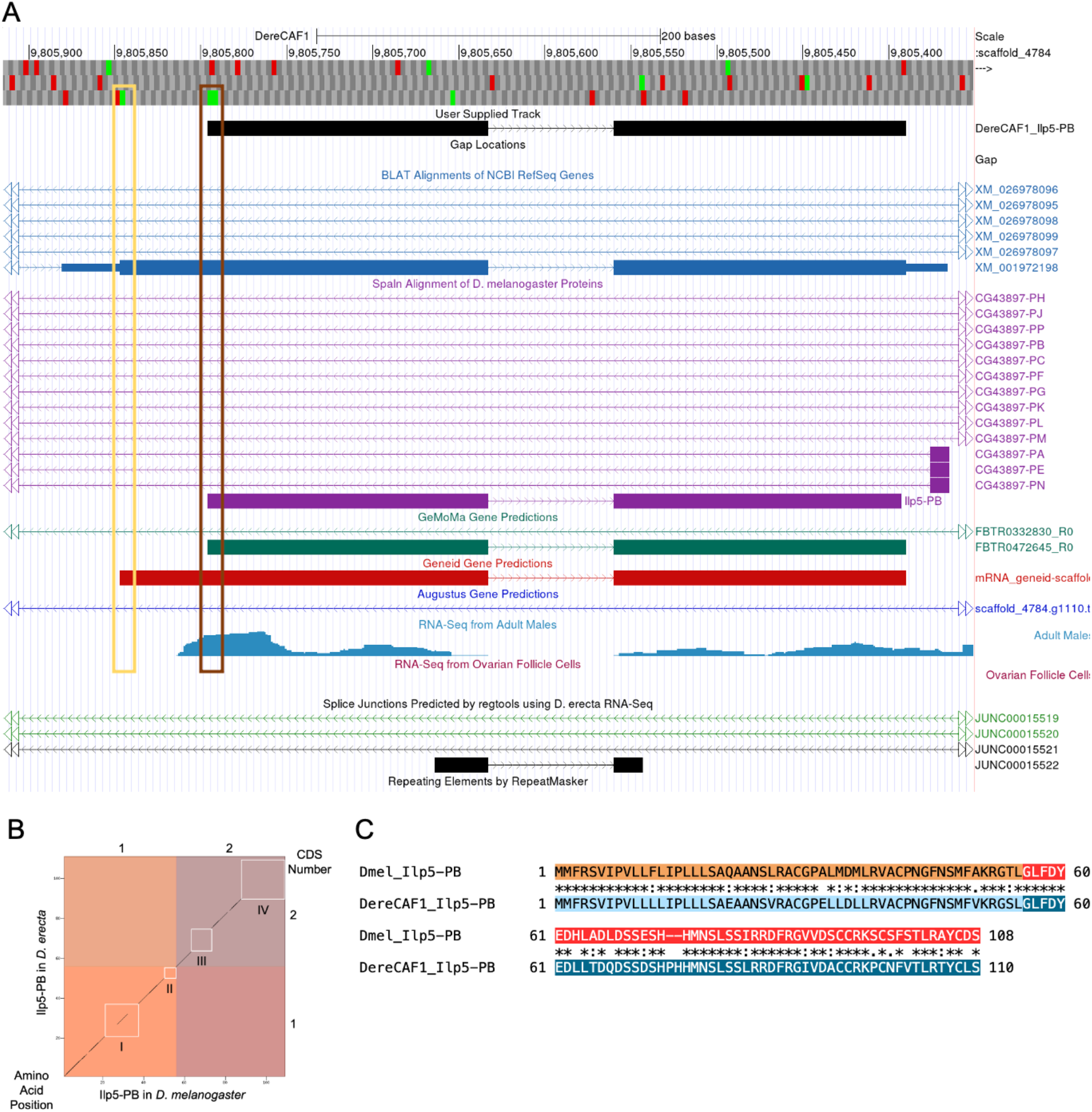
*D. erecta Ilp5* Gene Model. A. Gene Model in UCSC Track Data Hub on Dere/CAF. (Raney *et al*., 2014). The coding regions of *Ilp5* in *D. erecta* are displayed in the User Supplied Track (top black); CDSs are depicted by thick rectangles and introns are displayed as thin lines with arrows indicating the direction of transcription. Other evidence tracks include BLAT Alignments of NCBI RefSeq Genes (blue, alignment of Ref-Seq genes for *D. erecta*) with thin rectangles representing non-coding regions of exons and thick rectangles and arrows as above, Spaln alignment of *D. melanogaster* Proteins (purple, alignment of Ref-Seq proteins from *D. melanogaster*), GeMoMa and Geneid gene predictions (green and red, respectively), transcript abundance and location (blue for males) from RNA-seq data, the absence of RNA-Seq (magenta if present for females) from ovarian follicle cells, and splice junctions in black at bottom (JUNC00015522) has a read-depth of 1. The yellow and brown boxes indicate two potential start codons (see text). **B. Dot Plot of Ilp5-PB in *D. melanogaster* (*x*-axis) versus the orthologous polypeptide in *D. erecta* (*y*-axis)**. The dot plot was prepared with a word size of 3. Amino acid number is indicated along the left and bottom; coding sequences numbers are indicated along the top and right, and CDSs are also highlighted with alternating colors. The white boxes denoted I, II, III, and IV outline regions of the largest amino acid discrepancy between *D. melanogaster* and *D. erecta*. **C. Protein Alignment of Ilp5-PB in *D. melanogaster* (*top*) versus its ortholog in *D. erecta* (bottom)**. CDSs of *D. melanogaster* Ilp5-PB are shown in orange and red while the homologous CDSs of *D. erecta* Ilp5-PB are shown in light and darker blue. Asterisks indicate amino acid identity, colons and periods represent strong and weak amino acid similarity, respectively, and empty spaces indicate dissimilar amino acids. Dashes in the sequence indicate gaps in the protein relative to the other ortholog.

### Absence of Ovarian Follicle RNA-seq Coverage for the *Ilp5* Gene Model

The May 2011 (Agencourt dere_caf1/DereCAF1) assembly in *D. erecta* has RNA-Seq data for adult males and ovarian follicles (Figure 2A). The adult male RNA-Seq data support the proposed *Ilp5* gene model, whereas there is no RNA-Seq data seen for ovarian follicles. The Aug. 2014 (BDGP Release 6 + ISO1MT/dm6) assembly in *D. melanogaster* showed no RNA-Seq coverage in the embryonic development stage 0-8 hours, and insufficient RNA-Seq data in the embryonal development stage 8-24 hours within the modENCODE RNA-Seq (Developmental) (R5) track (Brown *et al*., 2014). Because there is no RNA-Seq coverage found within the first hours of embryonic development, it is expected that there is no RNA-Seq coverage found within the ovarian follicle stage. However, to verify this information, additional RNA-Seq data in form of long-RNA sequence data for ovarian follicle tissue in *D. melanogaster* is needed to confirm this information.

## Methods

“Detailed methods including algorithms, database versions, and citations for the complete annotation process can be found in Rele *et al*. (2023). Briefly, students use the GEP instance of the UCSC Genome Browser v.435 (https://gander.wustl.edu; Kent WJ *et al*., 2002; Navarro Gonzalez *et al*., 2021) to examine the genomic neighborhood of their reference IIS gene in the *D. melanogaster* genome assembly (Aug. 2014; BDGP Release 6 + ISO1 MT/dm6). Students then retrieve the protein sequence for the *D. melanogaster* reference gene for a given isoform and use *tblastn* against their target *Drosophila* species genome assembly on the NCBI BLAST server (https://blast.ncbi.nlm.nih.gov/Blast.cgi; Altschul *et al*., 1990) to identify potential orthologs. To validate the potential ortholog, students compare the local genomic neighborhood of their potential ortholog with the genomic neighborhood of their reference gene in *D. melanogaster*. This local synteny analysis includes at minimum the two upstream and downstream genes relative to their putative ortholog. They also explore other sets of genomic evidence using multiple alignment tracks in the Genome Browser, including BLAT alignments of RefSeq Genes, Spaln alignment of *D. melanogaster* proteins, multiple gene prediction tracks (e.g., GeMoMa, Geneid, Augustus), and modENCODE RNA-Seq from the target species. Detailed explanation of how these lines of genomic evidenced are leveraged by students in gene model development are described in Rele *et al*. (2023). Genomic structure information (e.g., CDSs, intron-exon number and boundaries, number of isoforms) for the *D. melanogaster* reference gene is retrieved through the Gene Record Finder (https://gander.wustl.edu/~wilson/dmelgenerecord/index.html; Rele *et al*., 2023). Approximate splice sites within the target gene are determined using *tblastn* using the CDSs from the *D. melanogaste*r reference gene. Coordinates of CDSs are then refined by examining aligned modENCODE RNA-Seq data, and by applying paradigms of molecular biology such as identifying canonical splice site sequences and ensuring the maintenance of an open reading frame across hypothesized splice sites. Students then confirm the biological validity of their target gene model using the Gene Model Checker (https://gander.wustl.edu/~wilson/dmelgenerecord/index.html; Rele *et al*., 2023), which compares the structure and translated sequence from their hypothesized target gene model against the *D. melanogaster* reference gene model. At least two independent models for a gene are generated by students under mentorship of their faculty course instructors. Those models are then reconciled by a third independent researcher mentored by the project leaders to produce the final model. Note: comparison of 5’ and 3’ UTR sequence information is not included in this GEP CURE protocol.” (Lawson *et al*., 2025). This gene model can also be seen within the target genome at this TrackHub link: D. erecta Ilp5 TrackHub.

## Supporting information

Gene model data files

## Supplemental Material

Zip file containing FASTA, PEP, and GFF files of the model

## Acknowledgments

We thank Wilson Leung, Washington University, St. Louis, for developing and maintaining the technological infrastructure that was used to create this gene model and Chinmay Rele and Laura K. Reed, University of Alabama, for their support and encouragement throughout this process. We are grateful to FlyBase for providing the definitive database for *Drosophila melanogaster* gene models.

